# Maintenance of spatial gene expression by Polycomb-mediated repression after formation of a vertebrate body plan

**DOI:** 10.1101/468769

**Authors:** Julien Rougeot, Naomi D. Chrispijn, Marco Aben, Dei M. Elurbe, Karolina M. Andralojc, Patrick J. Murphy, Pascal W.T.C. Jansen, Michiel Vermeulen, Bradley R. Cairns, Leonie M. Kamminga

## Abstract

Polycomb group (PcG) proteins are transcriptional repressors that are important regulators of cell fate during embryonic development. Among them, Ezh2 is responsible for catalyzing the epigenetic repressive mark H3K27me3 and is essential for animal development. The ability of zebrafish embryos lacking both maternal and zygotic *ezh2* to form a normal body plan provides a unique model to comprehensively study Ezh2 function during early development in vertebrates. By using a multi-omics approach, we found that Ezh2 is required for the deposition of H3K27me3 and is essential for the recruitment of Polycomb group protein Rnf2. However, and despite the complete absence of PcG-associated epigenetic mark and proteins, only minor changes in H3K4me3 deposition and gene and protein expression occurred. These changes were mainly due to local deregulation of transcription factors outside their normal expression boundaries. Altogether, our results in zebrafish show that Polycomb-mediated gene repression is important right after the body plan is formed to maintain spatially restricted expression profiles of transcription factors and highlight the differences that exist in the timing of PcG protein action between vertebrate species.

**Summary statement:** Our unique zebrafish model of maternal and zygotic mutant for the *Polycomb* group gene *ezh2* reveals major conserved and divergent mechanisms in epigenetic gene repression during vertebrate development.

## Introduction

Development of multi-cellular organisms involves highly dynamic and controlled processes during which one single totipotent cell will multiply and differentiate into all the cells composing the adult individual. Specification of cell identity is controlled through the establishment of spatially and temporally restricted transcriptional profiles, which are subsequently maintained by epigenetic mechanisms (Brock and Fisher, 2005). Epigenetic maintenance of gene expression can act through modifications of the chromatin, the complex of DNA wrapped around an octamer of histones H2A, H2B, H3, and H4 and its associated proteins and non-coding RNAs, creating an epigenetic landscape, often referred to as the epigenome (Zhu and Li, 2016). These modifications can be propagated from mother to daughter cells and thereby maintain gene expression profiles by controlling the accessibility of the DNA to the transcriptional machinery (Li and Reinberg, 2011).

Important regulators of the epigenome during development are the Polycomb Group (PcG) proteins. First identified in *Drosophila melanogaster,* PcG proteins were found to maintain the pre-established pattern of *hox* gene expression (Kennison, 1995). Subsequent studies showed that PcG proteins are important for proper patterning during early embryonic development, tissue-specific development, and maintenance of the balance between pluripotency and differentiation of stem cells in multiple species (Schuettengruber et al., 2017). Two main PcG complexes have been described (Chittock et al., 2017). The Polycomb Repressive Complex 2 (PRC2) is composed of the core subunits EZH1/2 (Enhancer of Zeste Homologue 1/2), SUZ12 (Supressor of Zeste 12), and EED (Embryonic Ectoderm Development). EZH2 has a catalytically active SET domain that trimethylates lysine 27 of histone H3 (H3K27me3), an epigenetic mark associated with gene repression and found along gene coding sequences (Mikkelsen et al., 2007). The catalytic subunits of PRC2 are mutually exclusive and EZH1 is postulated to complement the function of EZH2 in non-proliferative adult organs (Margueron et al., 2008, Shen et al., 2008). H3K27me3 can be recognized by the Polycomb Repressive Complex 1 (PRC1). A diversity of PRC1 compositions has been described and canonical PRC1 is composed of the core subunits RING1/RNF2 (Ring Finger Protein 2 a/b), PCGF1-6 (Polycomb Group RING fingers 1-6), PHC (Polyhomeotic), and CBX (Chromobox homolog) (Gao et al., 2012, Kloet et al., 2016). PRC1 catalyzes the ubiquitination of lysine 119 of histone H2A (H2AK1119ub) and strengthens gene repression. In contrast to this canonical view, recent studies implicate that PRC1 is also active in the absence of PRC2 (He et al., 2013, Loubiere et al., 2016, Tavares et al., 2012). Trithorax Group (TrxG) proteins antagonize PcG protein function through the deposition of a trimethyl group on lysine 4 of histone H3 (H3K4me3) on promoters and enhancers from virtually all transcribed genes (Klymenko and Muller, 2004, Schmitges et al., 2011, Santos-Rosa et al., 2002).

In mice, loss of PRC2 genes *Ezh2, EED,* or *Suz12* or PRC1 gene *Rnf2* leads to post-implantation embryonic lethality during early gastrulation (O’Carroll et al., 2001, Faust et al., 1998, Pasini et al., 2004, Voncken et al., 2003), making it difficult to study transcriptional regulation by PcG complexes during early development. Apart from the mouse model, very few studies have focused on functional characterization of PcG function during vertebrate development, and especially studies involving genetic mutants for different PcG genes. Lately, the zebrafish embryo has emerged as a model of choice to study developmental epigenetics in vertebrates (Chrispijn et al., 2019, Lindeman et al., 2011, Murphy et al., 2018, Potok et al., 2013, Vastenhouw et al., 2010). Others and we previously used loss-of-function mutants to show that *ezh2* is essential for zebrafish development (Dupret et al., 2017, San et al., 2016, San et al., 2018, Zhong et al., 2018). More particularly, our unique vertebrate model of zebrafish embryos mutant for both maternal and zygotic *ezh2,* referred as *MZezh2* mutant embryos, develop seemingly normal until 1 dpf, forming a proper body plan. These mutants ultimately die at 2 dpf, exhibiting a 100% penetrant pleiotropic phenotype associated with a loss of tissue maintenance (San et al., 2016). This makes zebrafish *MZezh2* mutant embryos a unique model to study the function of Ezh2 during early development, from fertilization to tissue specification, in the unique context of a vertebrate embryo in which trimethylation of H3K27 has never occurred, unlike cell culture, conditional, or zygotic mutant models.

We conducted a multi-omics approach in these *MZezh2* mutant embryos to study how PcG-mediated gene regulation controls axis formation and tissue specification. We focused our study on 24 hours post fertilization (hpf) embryos, when the first phenotypes become visible, and the anterior-posterior patterning of the embryos is properly established. Our results show conservation of basic PcG recruitment and silencing mechanisms and reveal that proper PRC2 function is essential for Rnf2 recruitment. However, very surprisingly, and despite the complete absence of PcG proteins and their associated epigenetic marks on the chromatin, the transcriptional and proteomic profile of *MZezh2* mutant embryos remains largely unchanged compared to wildtype embryos. The changes affect primarily a subset of PcG target genes. These genes are mainly transcription factors essential for developmental processes which present locally restricted aberrant gene expression. Our results show that zebrafish embryo development is initially independent of PcG repression until the stage of tissue maintenance and stress the differences that exist in the timing of PcG function requirement between vertebrate species.

## Results

### The repressive epigenetic mark H3K27me3 is absent in *MZezh2* embryos

To study the function of Ezh2 during development, we used the *ezh2* nonsense mutant allele *ezh2 (hu5670)* containing a premature stop codon within the catalytic SET domain, resulting in the absence of Ezh2 protein (San et al., 2016). Total elimination of both maternal and zygotic contribution of Ezh2 protein and RNA, by using the germ cell transplantation technique described previously (Ciruna et al., 2002, San et al., 2016), allowed us to study the function of Ezh2 during early development. As previously shown, *MZezh2* mutant embryos display normal body plan formation and a mild phenotype at 24 hpf. They die at 48 hpf, at which point pleiotropic phenotypes are observed, such as smaller eyes, smaller brain, blood coagulation, and absence of pectoral fins (Figure. 1A). Western Blot analysis at 3.3 hpf and 24 hpf confirmed the absence of both maternal and zygotic Ezh2 in these mutants, respectively (Figure 1B). In addition, our previous study also reported that H3K27me3 was not detectable in *MZezh2* mutants by immunofluorescence (San et al., 2016).

**Figure 1.**
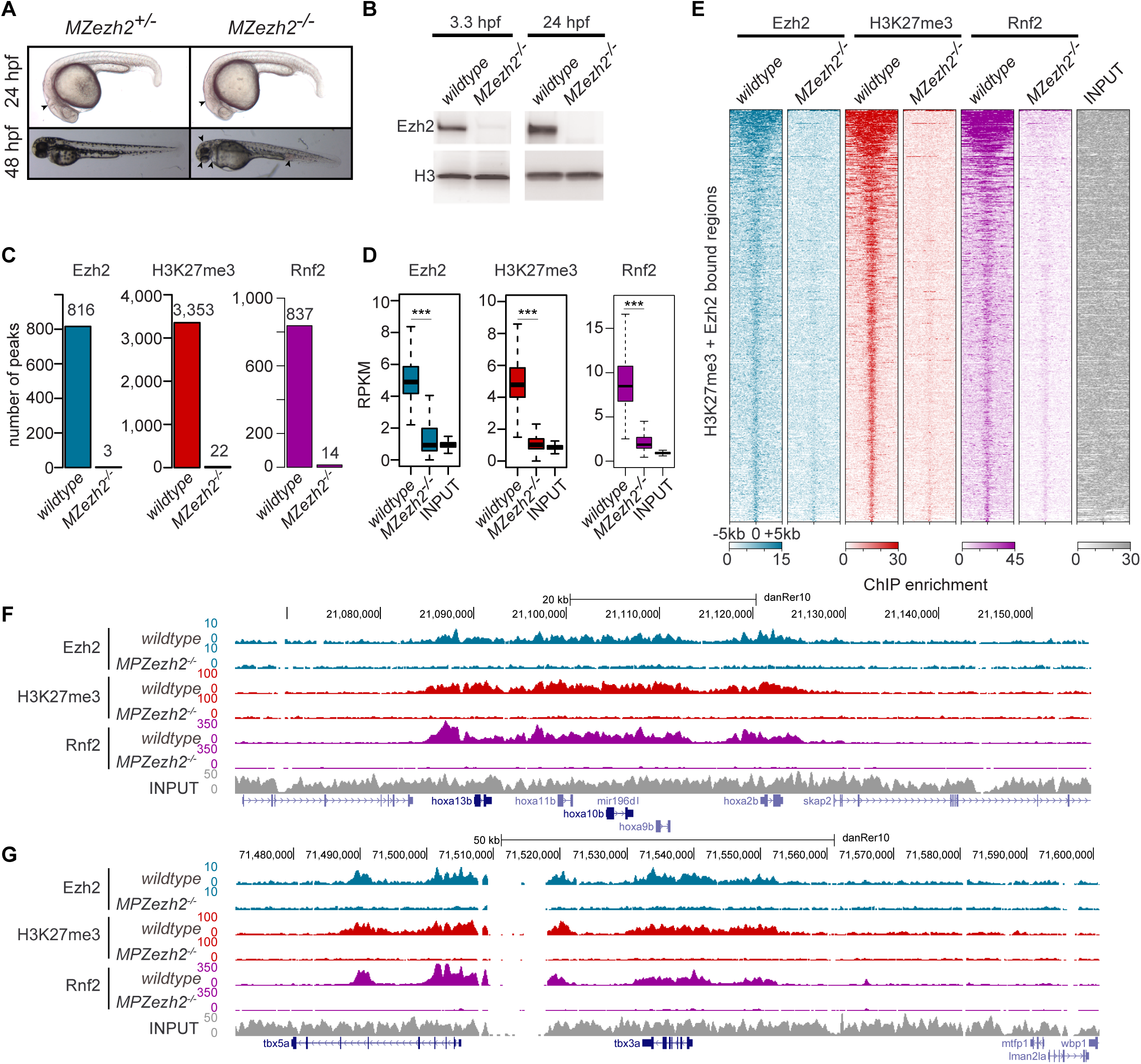
*MZezh2* mutant (*MZezh2*^*-/-*^) embryos lack Ezh2, H3K27me3, and Rnf2 binding to the chromatin. **(A)** *MZezh2*^*+/-*^ (developing as wildtype embryos) and *MZezh2*^*-/-*^ embryos at 24 and 48 hpf. At 24 hpf, *MZezh2*^*-/-*^ embryos lack a clear mid-hindbrain boundary compared to heterozygous embryos (arrow head). At 48 hpf, *MZezh2*^*-/-*^ embryos showed pleiotropic phenotypes compared to heterozygous embryos, such as small eyes, small brain, heart edema, and blood accumulation in the blood island (arrow heads). **(B)** Western blot analysis of Ezh2 at 3.3 hpf and 24 hpf of wildtype and *MZezh2*^*-/-*^ embryos. Histone H3 was used as a loading control. **(C)** Number of peaks called after Ezh2, H3K27me3, and Rnf2 ChIP-seq of wildtype and *MZezh2*^*-/-*^ embryos at 24hpf. Each peak set was obtained by the intersection of two independent biological replicates. **(D)** Box plots of Ezh2, H3K27me3, and Rnf2 RPKM-normalized coverage after respective ChIP-seq in wildtype and in *MZezh2*^*-/-*^ embryos at 24 hpf. The input control shown was obtained from wildtype embryos at 24 hpf. Coverages were calculated based on positions of peaks detected in wildtype embryos, t-test: *** *P*-value < 0.001. **(E)** Heatmaps for Ezh2, H3K27me3, and Rnf2 subsampled counts after ChIP-seq in 24 hpf wildtype and *MZezh2*^*-/-*^ embryos. Windows of 10 kb regions for all H3K27me3 o Ezh2 peaks in 24 hpf wildtype embryos are shown. The input track obtained from 24 hpf wildtype embryos was used as control and was not subsampled. **(F, G)** UCSC genome browser snapshot depicting the loss of Ezh2, H3K27me3, and Rnf2 after ChIP-seq in 24 hpf *MZezh2*^*-/-*^ embryos compared to wildtype embryos for **(F)** the *hoxab* gene cluster and **(G)** *tbx5a.* Colors represent ChIP-seq for different proteins with blue: Ezh2, red: H3K27me3, purple: Rnf2, and grey: Input control.

To further confirm the absence of Ezh2 in *MZezh2* mutants and its effect on H3K27me3 deposition, we performed ChIP-sequencing (ChIP-seq) for Ezh2 and H3K27me3 at 24 hpf in both wildtype and *MZezh2* mutant embryos. ChIP-seq analyses for Ezh2 and H3K27me3 revealed 816 and 3,353 peaks in wildtype embryos, respectively (Figure 1C, Table S1). Although the number of peaks differed between the two proteins, their binding profiles greatly overlap (Figure 1E). Quantification showed that 85% of Ezh2 peaks also contain H3K27me3 (Fig. S1A). Known PcG target genes such as the *hoxab* gene cluster, *tbx* genes, and *gsc* presented similar binding profiles for Ezh2 as for H3K27me3 (Figure 1F,G, Fig. S1B), whereas the ubiquitously expressed genes *eif1ad* and *tbp* showed absence of both Ezh2 and H3K27me3 (Fig. S1B).

In *MZezh2* mutant embryos, the binding of Ezh2 and H3K27me3, as detected by ChIP-seq, was virtually absent, with 3 and 22 peaks detected for Ezh2 and H3K27me3, respectively (Figure 1C). Manual inspection of these remaining peaks revealed that they are present in gene deserts and low complexity regions and are most probably artefacts (Fig. S1B). Ezh2 and H3K27me3 coverage was reduced to background levels in *MZezh2* mutants compared to wildtype (Figure 1D). Finally, the *hoxab* gene cluster, *tbx3a, tbx5a, gsc,* and *isl1* loci, targeted by PcG repression in wildtypes, also showed a complete absence of Ezh2 and H3K27me3 binding in *MZezh2* mutants (Figure 1F,G, Fig. S1B).

In order to rule out the possibility that the absence of detection of Ezh2 and H3k27me3 in *MZezh2* mutant samples was due to an inefficient ChIP-seq or a normalization artifact specific to mutant samples, the second ChIP-seq replicates for both Ezh2 and H3K27me3 were conducted with spike-in chromatin control. After normalization using the immunoprecipitated spike-in chromatin, the decrease in Ezh2 and H3K27me3 coverage in mutant compared to wildtype appear even higher than without normalization, both at genome-wide level (Fig. S2A,B) and on target genes (Fig. S2C).

Altogether, these results demonstrate that *MZezh2* mutants exhibit complete absence of Ezh2 and H3K27me3 from chromatin.

### Loss of PRC2-mediated repression results in loss of PRC1 recruitment during early development

It is postulated that PRC1 is recruited to chromatin by PRC2-deposited H3K27me3 but can also have a function independent of PRC2 (He et al., 2013, Loubiere et al., 2016, Tavares et al., 2012). As both Ezh2 and H3K27me3 are absent from *MZezh2* mutant embryos, we investigated whether PRC1 is still recruited to chromatin in these mutants. In zebrafish, Rnf2 is the only catalytic subunit of PRC1 (Le Faou et al., 2011). ChIP-seq for Rnf2 in wildtype embryos at 24 hpf reveals 837 peaks (Figure 1C, Table S1) which are present at Ezh2 and H3K27me3 positive regions (Figure 1E). We found that 70% of Ezh2 peaks were also positive for Rnf2 in wildtype embryos (Fig. S1A).

In *MZezh2* mutant embryos, only 14 binding sites could be detected for Rnf2 (Figure 1C) and Rnf2 average binding (measured in RPKM) was reduced to background level, as observed for Ezh2 and H3K27me3 binding (Figure 1C,E). This loss of Rnf2 was observed at both gene clusters such as *hoxab* (Figure 1F) and individual transcription factors such as *tbx3a, tbx5a,* and *gsc* (Figure 1G, Fig. S1B). As for Ezh2 and H3K27me3, Rnf2 remaining peaks in *MZezh2* mutant embryos were detected in intergenic regions with repeat sequences and are most probably artefacts (Fig. S1D).

Furthermore, H2AK119ub was not detectable in core histone extracts from *MZezh2* mutant embryos (Fig. S1B), suggesting a total absence of functional recruitment of canonical PRC1 in absence of Ezh2.

### Loss of H3K27me3 in *MZezh2* mutant embryos induces gene specific gain of H3K4me3

As PcG and TrxG complexes are known to have an antagonistic effect on gene expression (Piunti and Shilatifard, 2016), we investigated whether the loss of H3K27me3 in *MZezh2* mutant embryos changed the deposition H3K4me3, a mark associated with gene activation.

To this aim, we performed ChIP-seq for H3K4me3 in triplicates in both wildtype and *MZezh2* mutant embryos at 24 hpf. We observed a similar distribution of H3K4me3 peaks, with 10,556 peaks detected in wildtype embryos and 10,096 in *MZezh2* mutants (Figure 2A, Table S1). A majority of 9550 peaks were shared between wildtype and *MZezh2* mutant embryos (Figure 2A), suggesting little to no differences in H3K4me3 deposition in absence of Ezh2.

**Figure 2.**
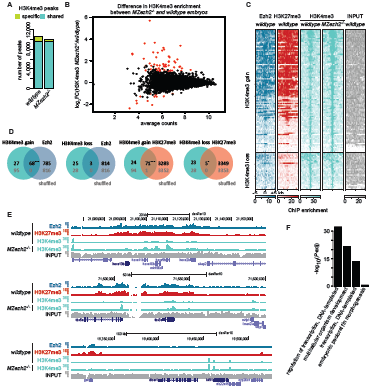
*MZezh2* mutant (*Mzezh2*^*-/-*^) embryos show an increase in H3K4me3 preferentially on H3K27me3 targets. **(A)** Number of peaks called after H3K4me3 ChIP-seq in wildtype and *Mzezh2*^*-/-*^ embryos at 24 hpf. Turquoise and green represent peaks shared by the two conditions and peaks specific for one condition, respectively. **(B)** MA-plot showing the fold change (log_2_-transformed) in H3K4me3 peak coverages in 24 hpf *MZezh2* mutant *(MZezh2*^*-/-*^) and wildtype embryos as a function of the normalized average count between the two conditions (log_10_-transformed) as calculated with DiffBind on the union of H3K4me3 peaks detected in both wildtype and *MZezh2* mutant conditions. Red: log_2_FC ≥ 1 or ≤-1 and *P*-adj < 0.05. **(C)** Heatmaps of subsampled counts after ChIP-seq for Ezh2 and H3K27me3 in wildtype embryos and H3K4me3 in wildtype and *Mzezh2*^*-/-*^ embryos at 24 hpf. Heatmaps display peaks enriched or decreased in *Mzezh2*^*-/-*^ embryos compared with wildtypes. The input track obtained from 24 hpf wildtype embryos was used as control. Blue, red, turquoise, and grey represent ChIP-seq for Ezh2, H3K27me3, H3K4me3, and Input control, respectively. **(D)** Venn diagrams presenting the overlap between peaks with increased or decreased H3K4me3 levels (gain or loss) as detected by DiffBind with the presence of Ezh2 or H3K27me3 peaks within a +/-1 kb window. Black numbers represent comparison between actual DiffBind results and ChIP-seq peaks whereas grey numbers represent comparisons between actual DiffBind results and randomly shuffled positions used as controls. X^2^: *** *P*-value **<** 0.001, **\*\*** *P*-value **<** 0.01, **\*** *P*-value **<** 0.05. **(E)** UCSC browser snapshots of three genomic loci in wildtype and *Mzezh2*^*-/-*^ embryos at 24 hpf. Blue, red, turquoise, and grey represent ChIP-seq for Ezh2, H3K27me3, H3K4me3, and Input control, respectively. **(F)** Gene Ontology analysis of the closest genes restricted two 2kb upstream or downstream from H3K4me3 peaks enriched in *Mzezh2*^*-/-*^.

We next assessed the differences in H3K4me3 peak intensity upon loss of Ezh2 by performing differential binding analysis using DiffBind. We identified 95 peaks with an enriched H3K4me3 deposition and 28 peaks with a decrease H3K4me3 intensity (Fig 2B). Comparison with Ezh2 and H3K27me3 ChIP-seq show an over-representation of PcG targets among the peaks enriched in H3K4me3 upon loss of Ezh2 (Fig 2C). A majority of the peaks enriched for H3K4me3 are PcG targets, with 71% (68 out of 95) targeted by Ezh2 and 75% (71 out of 95) by H3K27me3, which is more than expected by chance. On the opposite, peaks with a decrease of H3K4me3 deposition show little to no enrichment in PcG targets (Fig 2D). This result shows that the targets of PcG repression in wildtype are more susceptible to gain H3K4me3 upon loss of Ezh2/H3K27me3.

We then searched for the closest genes from the regions with increased H3K4me3 peak coverage detected by DiffBind and identified 118 genes. For example, the transcription factors *hoxa13b, tbx5a,* and *gsc* showed enrichment for H3K4me3 close to their promoter (Fig 2E). Gene ontology analysis revealed that these genes were mainly involved in transcriptional regulation and organismal development (Figure 2F). Among these 118 identified genes, 51 encode for transcription factors, among which were members of the *hox, tbx, sox,* and *pax* gene families, known targets of PcG complexes.

These results show that loss of PcG repression has a limited effect on H3K4me3 active epigenetic mark at 24 hpf, and that the genes presenting an increase in H3K4me3 deposition are mainly transcription factors directly targeted by PcG repression.

### Epigenetic changes in *MZezh2* mutant embryos induce upregulation of transcription factors

As *MZezh2* mutant embryos show a complete lack of the H3K27me3 repressive mark and a subtle yet selective increased deposition of H3K4me3 activating mark on genes coding for transcription factors, we investigated the effect of loss of Ezh2 on the transcriptome and proteome of wildtype and *MZezh2* mutant embryos at 24 hpf.

Transcriptome analysis by RNA-seq in the two conditions revealed a limited effect on the transcriptome upon the loss of Ezh2. Only 60 genes were detected to be significantly upregulated (log_2_FC ≥ 1 and *P*-adj < 0.05) and 28 genes downregulated (log_2_FC ≤ −1 and P-adj < 0.05) in *MZezh2* mutants (Figure 3A). The proteome analysis, identified 111 upregulated (log_2_FC ≥ 1.5 and *P-adj <* 0.05) and 110 downregulated (log_2_FC ≤ −1.5 and *P-adj* < 0.05) proteins in *MZezh2* mutants compared to wildtype controls.

**Figure 3.**
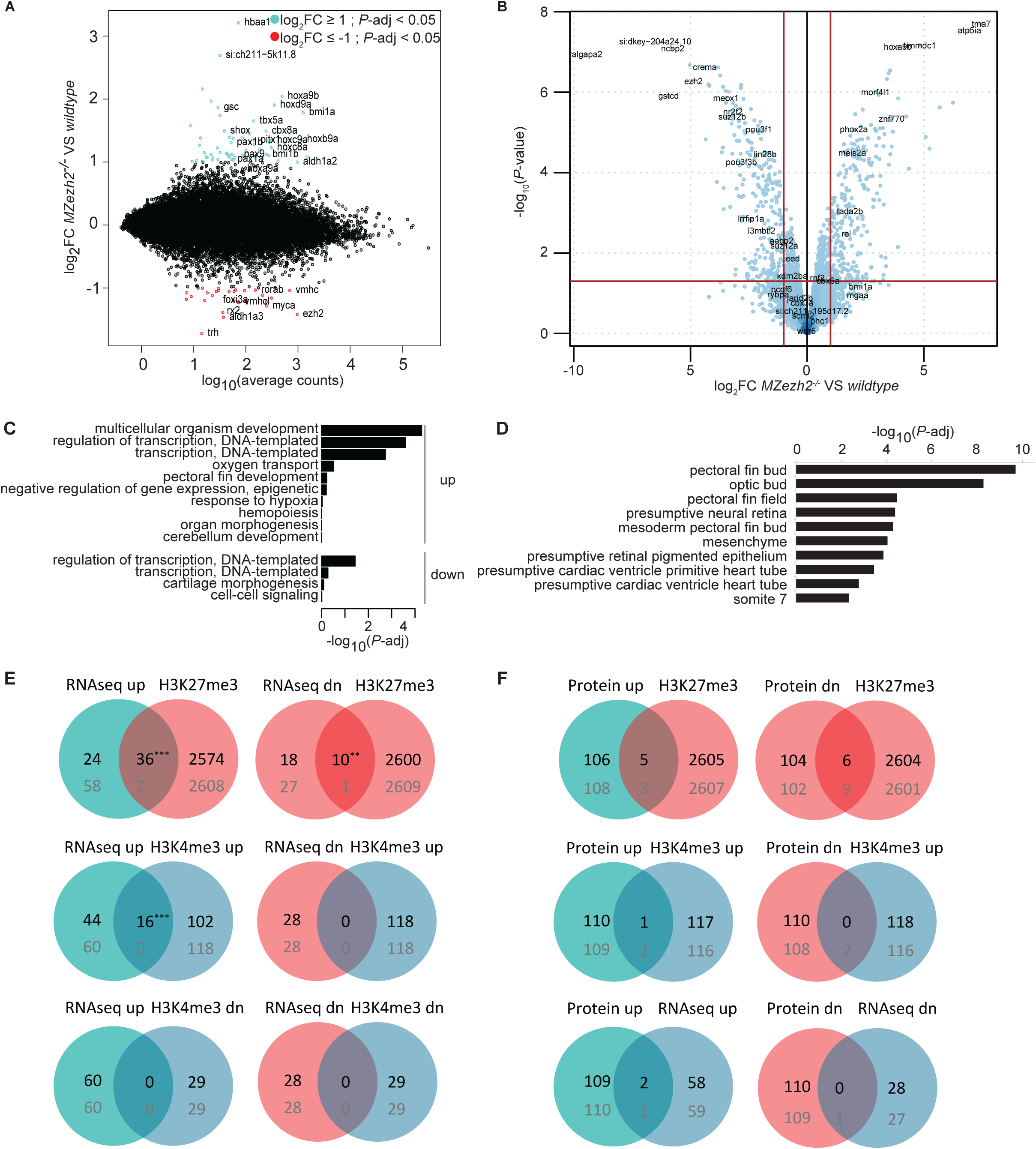
Loss of maternal zygotic *ezh2* results in overexpression of specific developmental genes. **(A)** MA-plot showing the fold change (log_2_-transformed) between gene expression in 24 hpf *MZezh2* mutant *(MZezh2*^*-/-*^) and wildtype embryos as a function of the normalized average count between the two conditions (log_10_-transformed) as calculated with DEseq2. Turquoise: log_2_FC ≥ 1 and *P*-adj < 0.05, red: log_2_FC < −1 and P-adj < 0.05. **(B)** Volcano plot showing the *P*-value (-log_10_-transformed) as a function of the fold-change (log_2_-transformed) between protein expression level in *MZezh2* mutant *(MZezh2*^*-/-*^) compared to wildtype embryos at 24 hpf. **(C)** Gene Ontology of biological processes analysis of genes upregulated (up) or downregulated (down) in *Mzezh2*^*-/-*^ embryos compared to wildtype embryos at 24 hpf. **(D)** Analysis of anatomical terms associated with genes upregulated and downregulated in *MZezhl*^*-/-*^ embryos compared to wildtype embryos at 24 hpf. **(E)** Venn diagrams presenting the overlap between genes upregulated (up) or downregulated (dn) in *MZezh2*^*-/-*^embryos compared to wildtype and presence of H3K27me3 or H3K4me3 peaks. The closest genes from H3K27me3 peaks in wildtype condition or H3K4me3 enriched (H3K4me3 up) and decreased (H3K4me3 dn) peaks according to DiffBind were used for this analysis. Black numbers represent comparison between actual DEseq2 identified genes and closest genes from peaks. Grey numbers represent comparisons between actual DEseq2 identified genes and random selected genes used as control. X^2^: *** *P*-value < 0.001, ** *P*-value < 0.01, * *P*-value < 0.05. **(F)** Venn diagrams presenting the overlap between proteins overrepresented (Protein up) or underrepresented (Protein dn) in *MZezh2*^*-/-*^ embryos compared to ChIP-seq and RNA-seq results. The closest genes from H3K27me3 peaks in wildtype condition or H3K4me3 enriched peaks according to DiffBind (H3K4me3 up) were used for this analysis. Black numbers represent comparison between actual deregulated proteins and genes. Grey numbers represent comparisons between actual deregulated proteins and random selected genes used as control. X^2^-testdid not provide any significant results.

GO analysis showed that the deregulated genes in the transcriptomics data are associated to control of organism development and regulation of transcription (Fig 3C). The proteins deregulated in the proteome analysis also revealed anatomy terms associated to organs presenting clear phenotypes in the *MZezh2* mutant embryos (Fig 3D).

When comparing our RNA-seq results with our ChIP-seq data, we found that 60% (36 out of 60) of the upregulated genes are targets of H3K27me3, which is more than expected by chance. Furthermore, a significant proportion (27%) of these genes also shows an increase in H3K4me3 deposition upon loss of Ezh2 whereas none show a decrease in H3K4me3 (Fig 3E). The downregulated genes also show some significant association with H3K27me3 but did not show any correlation with gain or loss of H3K4me3 deposition (Fig 3E). However, the proteomics data did not present any correlation with either the ChIP-seq or the RNA-seq results (Fig 3F).

Finally, proteome data indicate that, in addition to Ezh2, Suz12b is downregulated in *MZezh2* mutant embryos (Figure 4A). Other PRC2 subunits were either not detected or not significantly downregulated. Subunits of the canonical PRC1 complex were mostly not detected or not significantly overexpressed (Fig. S3).

**Figure 4.**
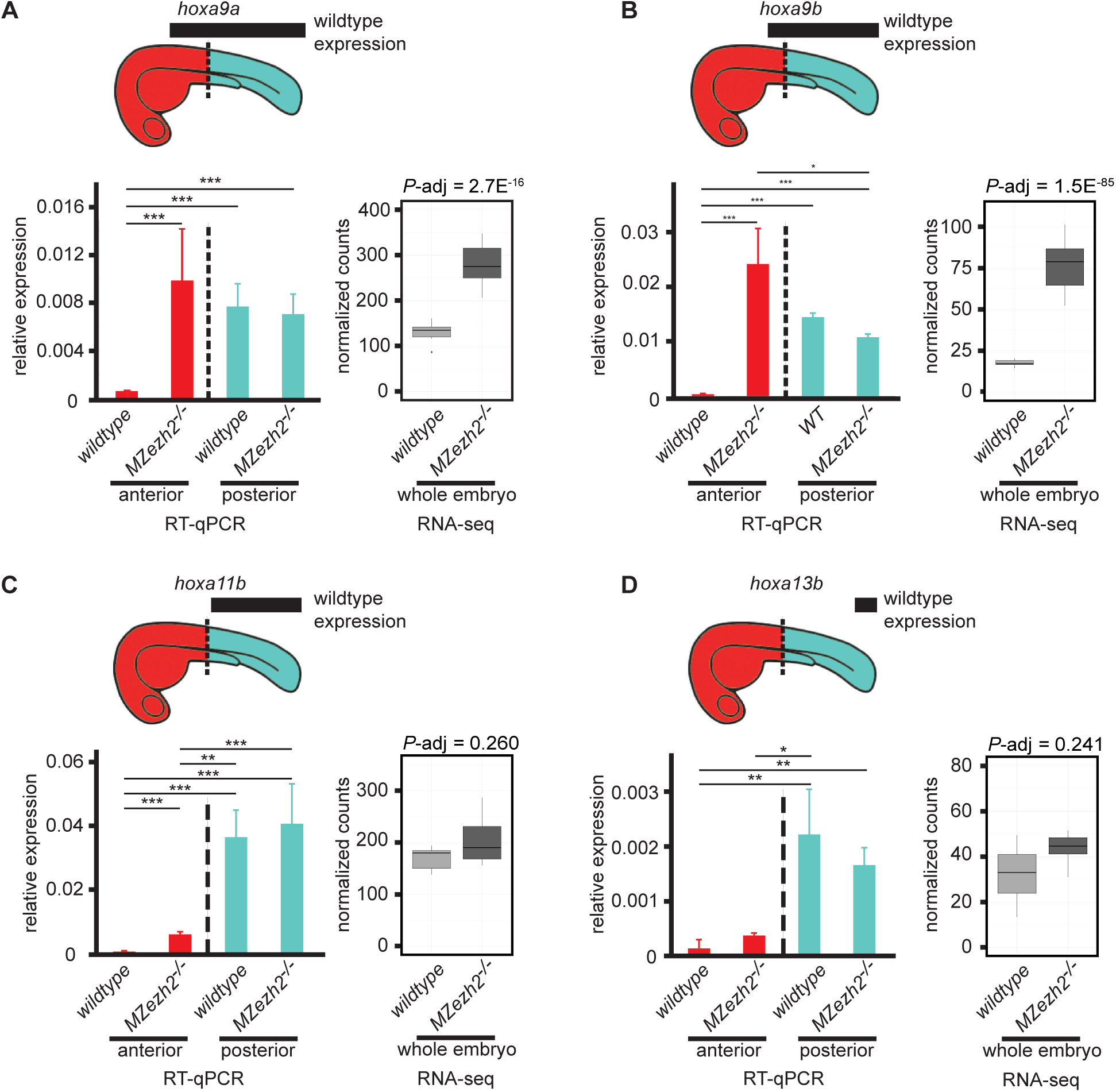
Loss of maternal and zygotic *ezh2* results in ectopic expression of *hox* genes, (a, b, c, d) Expression analysis of **(A)** *hoxa9a,* **(B)** *hoxa9b,* **(C)** *hoxa11b,* and **(D)** *hoxa13b* at 24 hpf. Bar plots on the left side of each panel represent relative expression of indicated *hox* genes in the anterior half (red) and posterior half (turquoise) of wildtype and *MZezh2* mutant (*MZezh2*^*-/-*^) embryos. Boxplots on the right side of each panel represent normalized counts from RNA-seq experiments in *MZezh2*^*-/-*^ and wildtype whole embryo lysates at 24 hpf. Above is a schematic representation of 1 dpf embryos. Black boxes represent the expression domains of the *hox* genes in wildtype embryos based on published data (Thisse, 2004). Dashed lines represent the demarcation between anterior (red) and posterior (turquoise) parts of the embryo used for RT-qPCR analysis. For RT-qPCR, relative expression was calculated based on expression of housekeeping gene *actb1.* Error bars represent standard deviations. Relative expression was compared between anterior or posterior parts in *MZezh2*^*-/-*^ and wildtype embryos (one-way ANOVA with post-tests, *** *P*-value < 0.001, ** *P*-value < 0.01, * *P-*value < 0.05). For RNA-seq, adjusted *P*-values were extracted from Differential Expression analysis with DEseq2.

### Loss of *ezh2* results in expression of *hox* genes outside their normal expression domains

We next carried out a spatial expression analysis on selected target genes to distinguish between the possibilities that absence of PcG-mediated repression leads to global but moderate gene deregulation or to deregulation limited to specific cell types or tissues.

To start with, we focused on expression of different genes from the *hox* gene family. These genes are known targets of Polycomb-mediated repression (Mallo and Alonso, 2013) and some of them have been previously shown to be deregulated in *MZezh2* mutant embryos (San et al., 2016). Every *hox* gene has an expression pattern that is restricted along the anterior-posterior axis (Prince et al., 1998). To obtain spatially resolved data along the anterior-posterior axis, we performed RT-qPCR on the anterior half and the posterior half of 24 hpf wildtype and *MZezh2* mutant embryos. We then compared the normalized relative expression levels between the different halves of the *MZezh2* mutant and wildtype embryos. The tested *hox* genes were selected based on their domain of expression along the anterior-posterior axis (Figure 5A-D). The *hoxa9a* gene, whose expression extends to anterior, until slightly outside the posterior half of the embryos, showed, as expected, a higher expression in the posterior part than in the anterior part in wildtype embryos (Figure 5A). In *MZezh2* mutants, *hoxa9a* was overexpressed in the anterior compartment compared with wildtype embryos, reaching levels similar to those observed in the posterior part of wildtype embryos. No significant differences were detected in the level of expression when comparing the posterior compartment of *MZezh2* mutant and wildtype embryos (Figure 5A). Similar results were obtained for *hoxa9b,* where overexpression was detected in the anterior compartment of *MZezh2* mutant embryos compared to the anterior compartment of wildtype embryos (Figure 5B). The *hoxa11b* and *hoxa13b* genes, which are expressed primarily posterior, showed, as expected, higher expression in the posterior half of the wildtype embryos compared to the anterior half (Figure 5C,D). In the *MZezh2* mutant embryos, both *hox* genes were upregulated in the anterior half of the *MZezh2* mutant embryos compared to wildtypes (Figure 5C,D) but their expression level remained lower than in the posterior half of the wildtype embryos (Figure 5C,D).

**Figure 5.**
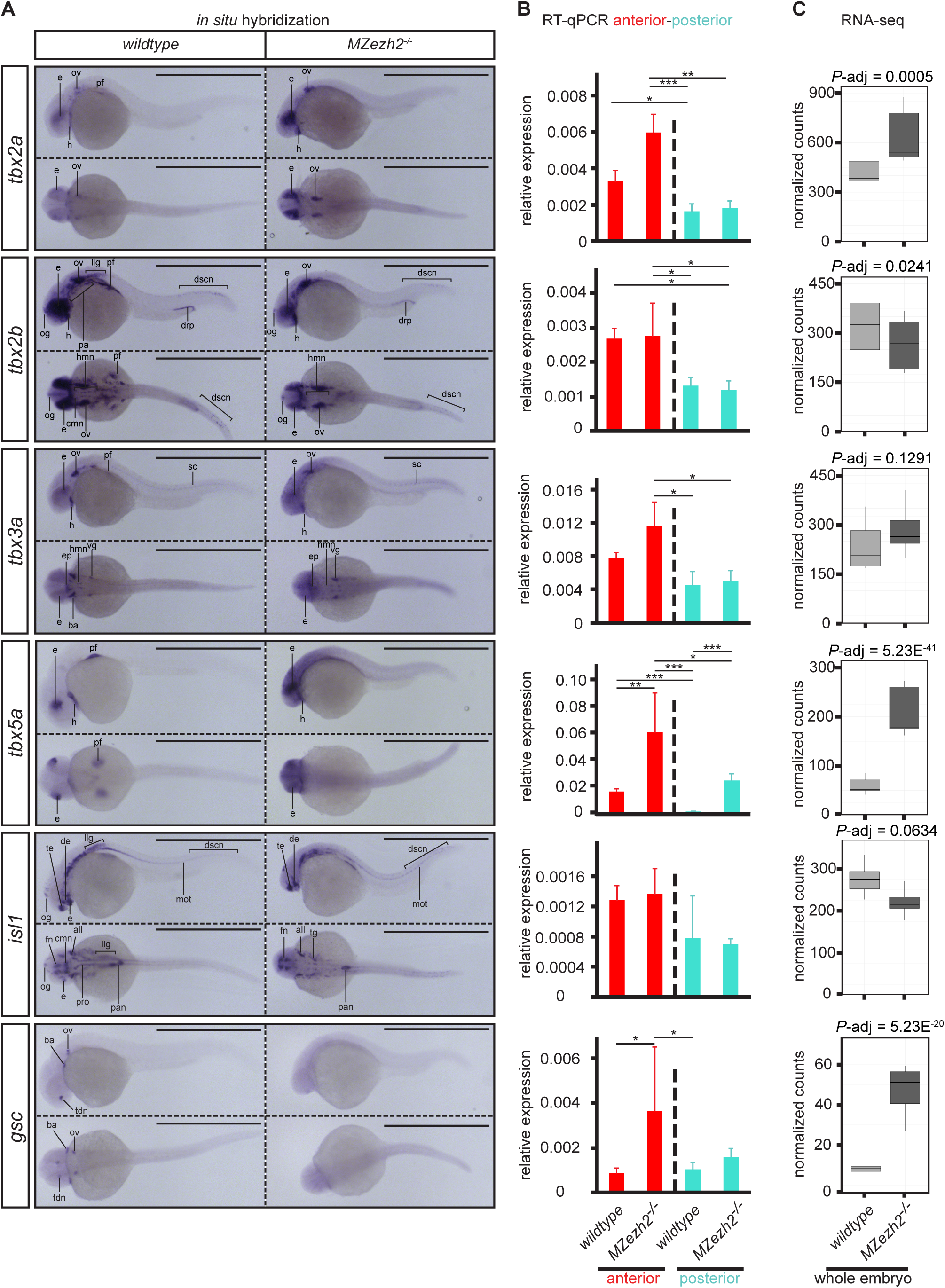
Transcription factor expression is spatially deregulated in *MZezh2* mutant (*MZezh2*^*-/-*^) embryos. **(A, B, C)** Spatial expression analysis by **(A)** *in situ* hybridization (ISH), **(B)** RT-qPCR on anterior half and posterior half, and **(C)** RNA-seq results of transcription factors *tbx2a, tbx2b, tbx3a, tbx5a, isl1,* and *gsc* in 24 hpf embryos. In ISH, scale bars represent 1 mm. For RT-qPCR, relative expression was calculated based on expression of housekeeping gene *actb1.* Error bars represent standard deviations. Relative expression was compared between anterior (red) or posterior (turquoise) parts in *Mzezh2*^*-/-*^ and wildtype embryos (one-way ANOVA with post-tests, *** *P*-value < 0.001, ** *P*-value < 0.01, * *P*-value < 0.05). Right boxplots represent normalized counts from RNA-seq experiments in whole *Mzezh2*^*-/-*^ and wildtype embryos and adjusted *P*-values were taken from Differential Expression analysis with DEseq2. all: anterior lateral lane ganglion, ba: branchial arch, cmn: cranial motor neurons, de: diencephalon, drp: distal region of the pronephros, dscn: dorsal spinal cord neurons, e: eye, ep: epiphysis, fn: forebrain nuclei, h: heart, hmn: hindbrain motor neurons, llg: lateral lane ganglion, mot: primary motor neurons, og: olfactory ganglion, ov: otic vesicle, pa: pharyngeal arches, pan: pancreas, pf: pectoral fin, pro: pronephros, sc: spinal cord, tdn: telencephalon and diencephalon nuclei, te: telencephalon, vg: ventral ganglion.

These comparative analyses of anterior and posterior parts of the embryo suggest that, upon loss of Ezh2, *hox* genes show an ectopic anterior expression while keeping wildtype expression levels in their normal expression domains.

### Different transcription factors show various profiles of deregulation in the absence of Ezh2

To further pursue our investigation on the changes in gene expression patterns in absence of Ezh2, we performed *in situ* hybridization (ISH) on members from the *tbx* gene family of transcription factors. The *tbx2a, tbx2b, tbx3a,* and *tbx5a* genes have partial overlapping expression patterns in wildtype embryos, but also display gene specific expression domains (Figure 6A). At 24 hpf, these *tbx* gene family members are expressed in the dorsal region of the retina, in the heart, and the pectoral fins (Ribeiro et al., 2007, Tamura et al., 1999). In addition, the genes *tbx2a, tbx2b,* and *tbx3a* are expressed in the otic vesicle. The genes *tbx2b* and *tbx3a* are expressed in different ganglions and neurons in anterior and posterior regions of wildtype embryos (Ribeiro et al., 2007). Finally, expression of *tbx2b* can also be detected in part of pharyngeal arches 3-7 and the distal region of the pronephros and *tbx3a* expression can be detected in the brachial arches (Thisse, 2004). This spatial prevalence of *tbx* gene expression in the anterior half of the embryo was also detected by RT-qPCR at 24 hpf, where *tbx2a, tbx2b* and *tbx5a* expression was significantly higher in the anterior than in the posterior part of wildtype embryos (Figure 6B).

ISH for these *tbx genes* on *MZezh2* mutant embryos at 24 hpf suggests ectopic expression of these transcription factors around their normal expression pattern in the eye, the otic vesicle, and the heart, except for *tbx2b* (Figure 6A). This scattering in gene expression was reflected in a trend towards a higher expression in the anterior half of *MZezh2* mutant embryos as detected by RT-qPCR, although only *tbx2a* and *tbx5a* results were significant (Figure 6B). In addition, ISH for *tbx5a,* and to a lesser extent *tbx3a,* showed ubiquitous expression throughout the entire body of *MZezh2* mutants which was not visible in wildtypes (Figure 6A). RT-qPCR results confirmed increased expression of *tbx5a* in both the anterior and posterior half of the *MZezh2* mutant embryos (Figure 6B).

Beside the observed ectopic expression, all tested *tbx* genes showed absence of expression in specific structures upon Ezh2 loss. For example, in *MZezh2* mutant embryos, all four *tbx* genes showed no expression in the fin bud (Figure 6A). In *MZezh2* mutant embryos, the gene *tbx2b* showed no expression in the pharyngeal arches 3-7 and the lateral line ganglions, and *tbx3a* was not observed in the branchial arches (Fig 6a). This absence of expression was not detected by RT-qPCR (Figure 6B) but a trend toward downregulation for *tbx2b* was observed in RNA-seq results on whole *MZezh2* mutant embryo lysates (Figure 6C).

In addition, we tested transcription factors from other families which were targeted by H3K27me3 in wildtype. The transcription factor *isl1,* expressed in all primary neurons (Dyer et al., 2014), showed a similar absence of expression in the fin bud and the cranial motor neurons in the midbrain (trigeminal, facial and vagal motor neurons), as observed for *tbx2a.* Its expression was also absent in the ventral region of the eye, the facial ganglia, and in the pronephros from *MZezh2* mutant embryos, where it is normally expressed in wildtype embryos (Heisenberg et al., 1999, Zhang et al., 2017) (Figure 6A). This loss in expression in *MZezh2* mutant embryos was not detected by RT-qPCR but a clear tendency toward downregulation was detected by RNA-seq (Figure 6B,C). Even more surprising was the expression pattern of *gsc* in the *MZezh2* mutant embryos. Whereas all the wildtype embryos show highly specific expression in the telencephalon and diencephalon nuclei, the branchial arches, and the otic vesicle (Thisse, 2004), *gsc* expression was lost and instead ubiquitous expression was observed in *MZezh2* mutant embryos (Figure 6A). This observation was confirmed by RT-qPCR and RNA-seq where upregulation of *gsc* was clearly detected in *MZezh2* mutant embryos (Figure 6B,C).

Taken together, these spatial expression analyses showed that the tested transcription factors are expressed outside their normal wildtype expression boundaries in *MZezh2* mutant embryos at 24 hpf. Furthermore, expression of some of these genes is lost in specific tissues in the *MZezh2* mutant embryos.

## Discussion

Here, we showed for the first time the genome-wide binding patterns of Ezh2 and Rnf2, the catalytic subunits of PRC2 and PRC1, respectively, in 24 hpf zebrafish embryos. The overall overlap between the binding patterns of Ezh2, Rnf2, and the PcG related epigenetic mark H3K27me3 suggests that the PcG-mediated gene repression mechanisms (Chittock et al., 2017) are evolutionary conserved in zebrafish development. The complete loss of H3K27me3 in *MZezh2* mutant embryos reveals that Ezh2 is the only methyltransferase involved in trimethylation of H3K27 during early zebrafish development. This result was expected as Ezh1, the only other H3K27me3 methyltransferase, was shown by a number of studies not to be maternally loaded nor expressed in the zebrafish embryo until at least after 1 dpf (Chrispijn et al., 2018, San et al., 2016, Sun et al., 2008, White et al., 2017). In addition, proteomics results showed decreased protein expression of most PRC2 subunits. This could indicate a destabilization of PRC2 in absence of the catalytic subunit in *MZezh2* mutant embryos. We could therefore confirm that zebrafish embryos can form a normal body plan in absence of PRC2-mediated gene repression.

The total loss of Rnf2 binding in the *MZezh2* mutants suggests that only the canonical pathway, in which PRC2 is required for PRC1 recruitment, is active during this stage of development. This absence of PRC1 recruitment to the chromatin is not caused by an absence of the complex in the *MZezh2* mutants, since most of the PRC1 subunits were detectable and not deregulated as shown by proteomic analysis. This is in contrast with studies in cultured mouse embryonic stem cells where non-canonical PRC1 complexes were shown to be recruited to developmental regulated genes independently of PRC2 (He et al., 2013, Tavares et al., 2012). This difference could be explained by the complete absence of H3K27me3 as from fertilization onwards in *MZezh2* mutant embryos, whereas other studies used conditional knockdown. Therefore, our model suggests that the PRC2-independent recruitment of PRC1 during early development can occur if PRC1 recruitment was first primed by a PRC2-dependent mechanism happening earlier during development.

As repressive and activating marks are known to antagonize each other (Schmitges et al., 2011), one could expect an increase in the H3K4me3 level deposited by TrxG proteins in absence of H3K27me3 associated with an increase in gene activation. However, the effects on H3K4me3 deposition, gene expression, and protein expression are limited in *MZezh2* mutant embryos at 24 hpf. This observation is in agreement with the near complete absence of phenotype at this developmental time point. Thus, it appears that transcriptional regulation during zebrafish development is largely PRC2-independent until later stages of development, when maintenance of cellular identity is required. These results were unexpected, as PRC2 is described to be essential during mammalian development already during gastrulation (Faust et al., 1998, O’Carroll et al., 2001, Pasini et al., 2004). It implies, that even if PcG-mediated repression mechanisms are conserved, the developmental stages at which these mechanisms are required differ between species. Possibly, the external development of the zebrafish and its rapid early development could explain this difference in phenotype.

Although limited, genes that show a gain in H3K4me3 deposition or in expression upon loss of *ehz2* are mainly transcription factors targeted by H3K27me3 in wildtype embryos. That only a minor fraction of all H3K27me3 target genes gained expression (36 out of 2610 = 1.2%, Fig 3E) suggests different mechanisms of regulation of PcG target genes at this time. Our hypothesis is that control of gene expression by signaling pathways and transcription factor networks (McGinnis and Tickle, 2005) is a robust mechanism and can be maintained until 1 dpf in absence of repression by PcG complexes. At 1 dpf, in absence of PcG-mediated repression, the first derepressed genes will be the genes subjected to the most fine-tuned transcriptional control, such as genes controlled by precise morphogen gradients. For example, it was shown that PRC2 attenuates expression of genes controlled by retinoic acid signaling (Laursen et al., 2013, Zhang et al., 2014). In vertebrates, and most particularly zebrafish, retinoic acid signaling is responsible for induction of formation of, among others, the forelimb field (Cunningham et al., 2013, Grandel and Brand, 2011), dorsoventral patterning of eyes (Lupo et al., 2005, Marsh-Armstrong et al., 1994), hindbrain patterning (Maves and Kimmel, 2005), *hox* gene expression (White et al., 2007), and the development of other organs (Samarut et al., 2015). All these processes are affected in *MZezh2* mutant embryos at 24 hpf and onwards and, therefore, could be explained by a defect in the response to retinoic acid signaling.

Spatial analysis of gene expression revealed different effects on gene expression patterns caused by loss of Ezh2. Anterior-posterior specific RT-qPCR showed that *hox* genes become abnormally expressed in the anterior half of the *MZezh2* mutant embryos; whereas expression levels in the posterior half remained unchanged. These results are in agreement with previous studies showing ectopic expression of *hox* genes in PRC1 and PRC2 zebrafish mutants (San et al., 2016, van der Velden et al., 2012), but also in other animal models (Kennison, 1995). Other transcription factors, such as the *tbx* gene family members, showed more diverse patterns of deregulation compared to *hox* genes. ISH and RT-qPCR showed that, among the *tbx* genes examined, some were overexpressed outside their normal expression domains *(tbx2a, tbx3a, and tbx5a),* whereas others were also ubiquitously upregulated *(tbx3a* and *tbx5a).* The case of eye patterning is a good example of the defect in control of gene expression pattern in *MZezh2* mutant embryos. In wildtype embryos, at 24 hpf, *tbx* genes are expressed in the dorsal part of the eye whereas *isl1* is expressed in the ventral part. Upon loss of Ezh2, our ISH results showed that the expression of the *tbx* genes expands to the whole eye whereas *isl1* disappears from the ventral region. We conclude that Polycomb-mediated repression is therefore responsible for maintenance of expression domains rather than control of expression level at this time of development in the zebrafish embryo.

Expression analysis by ISH for *tbx* and *hox* genes as well as for *isl1* also showed loss of expression in specific structures in *MZezh2* mutant embryos. We reasoned that the absence of expression of *hox* and *tbx* genes in the fin bud could be a secondary effect due to the absence of this structure in *MZezh2* mutants (San et al., 2016). The same phenomenon could explain the lack of detection of *tbx2b* and *isl1* in pharyngeal arches, pronephros, and lateral line ganglions. The case of *gsc* expression is more striking, as its normal expression pattern is totally abolished and a ubiquitous expression pattern is detected. The *gsc* gene is known to be expressed in the Spemann organizer during gastrulation and therefore all cells will transiently express *gsc* when undergoing gastrulation (Joubin and Stern, 1999, Stachel et al., 1993). In absence of Ezh2, *gsc* expression could remain active in all cells after leaving the Spemann organizer, leading to a ubiquitous expression pattern and impaired tissue specific expression in 24 hpf *MZezh2* mutant embryos.

To conclude, our results show that major characteristics of PcG-mediated repression are conserved in zebrafish, including canonical recruitment or PcG complexes and their function in maintenance of pre-established gene expression patterns. Our use of a mutant depleted of both maternal and zygotic contribution of Ezh2 also reveals that no PRC2-independent recruitment of PRC1 occurs at this stage of development. Finally, we demonstrate that early embryonic development, including germ layer formation and cell fate specification, is independent of PcG-mediated gene repression until axis are formed and organs specified. PcG-mediated gene repression is then required to control precise spatial restricted expression of specific transcription factors. We hypothesize that subtle changes in expression of these important genes subsequently will lead to progressive and accumulating changes in gene network regulation and result in loss of tissue identity maintenance.

This surprising result highlights the fact that, despite the conservation of PcG-mediated repression mechanisms during evolution, the time frame within which PcG repression is required for proper development may vary greatly between species. Studying the PcG repression in additional species would improve our understanding of the importance of PcG biology during development.

## Materials and methods

### Zebrafish genetics and strains

Zebrafish *(Danio rerio),* were housed according to standard conditions (Westerfield, 2000) and staged according to Kimmel et al. (Kimmel et al., 1995). The *ezh2* nonsense mutant *(hu5670)* (San et al., 2016), *Tg (H2A::GFP)* (Pauls et al., 2001), and *Tg (vas::eGFP)* (Krovel and Olsen, 2002) zebrafish lines have been described before. Genotyping of the *ezh2* allele was performed as previously described (San et al., 2016) with following adaptations: different primer pairs were used for PCR and nested PCR (Table S2), of which the restriction profile is shown on Fig. S1F. All experiments were carried out in accordance with animal welfare laws, guidelines, and policies and were approved by the Radboud University Animal Experiments Committee.

### Germ cell transplantation

Germ cell transplantation was performed as described previously (San et al., 2016). For all experiments below, *ezh2* germline mutant females were crossed with *ezh2* germline mutant males to obtain 100% *MZezh2* mutant progeny. The germline wild-type sibling males and females obtained during transplantation were used to obtain 100% wildtype progeny with similar genetic background and are referred to as wildtype. The embryos used were all from the first generation after germline transplantation.

### Western blotting

At 3.3 hpf, 50 embryos were collected, resuspended in in 500 μl ½ Ringer solution (55 mM NaCl, 1.8 mM KCl, 1.25 mM NaHCO_3_) and forced through a 21G needle and a cell strainer in order to remove the chorion and disrupt the yolk. At 24 hpf, 20 embryos were collected and resuspended by thorough pipetting in 500μl ½ Ringer solution in order to disrupt the yolk. The samples of 3.3 and 24 hpf were centrifuged for 5 minutes at 3,500 g at 4°C and washed two additional times with 500 μl ½ Ringer solution. The embryo pellet was frozen in liquid nitrogen and stored at −80°C. Whole protein extraction was performed by adding 40 μl of RIPA buffer (100 mM Tris-HCl pH 8, 300 mM NaCl, 2% NP-40, 1% Sodium Deoxycholate, 0.2% SDS, 20% glycerol, 1x cOmplete EDTA-free protease inhibitor cocktails from Sigma) and sonication for 2 cycles of 15s ON and 15s OFF on medium power at 4°C on a PicoBioruptor (Diagenode). After 10 minutes incubation at 4°C, embryo lysates were centrifuged for 12 minutes at 16,000 g at 4°C and supernatant was transferred in a new tube. 20 μg protein was mixed with SDS containing sample loading buffer, denaturated at 95°C for 5 minutes and analyzed by Western blot analysis. Antibodies used for immunoblotting are described in Table S3 HRP-conjugated anti-rabbit secondary antibody was used (Table S3) and protein detection was performed with ECL Select Western Blotting Detection Reagent (GE Healthcare, RPN2235) on an ImageQuant LAS 4000 (GE Healthcare) Western-blot anti-H2A western-blot were performed on histone extract according to van der Velden et al. (van der Velden et al., 2012), and detected on X-ray film.

### ChIP-sequencing

For chromatin preparation, embryos from a germline mutant or germline wildtype incross were collected at 24 hpf and processed per batches of 300 embryos. Embryos were first dechorionated by pronase (0.6 μg/μl) treatment and then extensively washed with E3 medium. Subsequently, embryos were fixed in 1% PFA (EMS, 15710) for 15 minutes at room temperature and fixation was terminated by adding 0.125M glycine and washed 3 times in cold PBS. Yolk from fixed embryos was disrupted by pipetting the fixed embryos 10 times with a 1 ml tip in 600 μl of ½ Ringer solution (55 mM NaCl, 1.8 mM KCl, 1.25 mM NaHCO_3_) and incubated for 5 minutes at 4°C on a rotating wheel. Embryos were pelleted by centrifuging 30 seconds at 300 g and the supernatant was removed. De-yolked embryos were resuspended in 600 μl sonication buffer (20 mM Tris-HCl pH 7.5, 70 mM KCl, 1 mM EDTA, 10% glycerol, 0.125% NP40, 1x cOmplete EDTA-free protease inhibitor cocktails from Sigma) and homogenized with a Dounce homogenizer (6 strokes with pestle A, followed by 6 strokes with pestle B). Homogenates were sonicated for 6 cycles of 30 seconds ON/30 seconds OFF on a PicoBioruptor (Diagenode), centrifuged for 10 minutes at 16,000 g at 4°C, and the supernatant containing the chromatin was stored at −80°C. 20 μl of the supernatant was subjected to phenol-chloroform extraction and ran on an agarose gel to verify that a proper chromatin size of 200-400 bp was obtained.

For ChIP, 100 μl of chromatin preparation (corresponding to 50 embryos) was mixed with 100 μl IP-buffer (50 mM Tris-HCL pH 7.5, 100 mM NaCl, 2 mM EDTA, 1% NP-40, 1x cOmplete EDTA-free protease inhibitor cocktails from Sigma) and antibody (for details on antibodies used see Table S3) and incubated overnight at 4°C on a rotating wheel. When relevant, *Drosophila* chromatin and anti-H2Av were used according to manufacturer’s instructions were followed (Active Motif, 53093 and 61686). For immunoprecipitation, 20 μl of protein G magnetic beads (Invitrogen, 1003D) were washed in IP buffer and then incubated with the chromatin mix for 2 hours at 4°C on a rotating wheel. Samples were washed in 500 μl washing buffer 1 (IP-buffer + 0.1% Sodium Deoxycholate), followed by washing in washing buffer 2 (washing buffer 1 + 400mM NaCl), washing buffer 3 (washing buffer 1 + 250mM LiCl), washing buffer 1 and a final wash in 250 μl of TE buffer. All washes were 5 minutes at 4°C on a rotating wheel. Chromatin was eluted from the beads by incubation in 100 μl of elution buffer (50 mM NaHCO_3_ pH 8.8, 1% SDS) for 15 minutes at 65°C at 900 rpm in a thermomixer. The supernatant was transferred in a clean 1.5 ml tube. Elution was repeated a second time and both supernatants were pooled. The eluate was treated with 0.33 μg/μl RNaseA for 2 hours at 37°C. Samples were then decrosslinked by adding 10 μl of 4M NaCL and 1 μl of 10mg/ml proteinase K and incubated overnight at 65°C. DNA was then purified using MinElute Reaction Clean-Up kit (Qiagen, 28204).

1-5 ng of DNA was used to prepare libraries with the KAPA Hyper Prep Kit (KAPABiosystems, KK8504) and NEXTflex ChIP-Seq Barcodes for Illumina (Bioo Scientific, 514122) followed by paired-end 43bp sequencing on an Illumina NextSeq500 platform. All ChIP-seq were performed in two biological replicates, except for Rnf2 in wildtype embryos which was performed once and H3K27ac which was performed in triplicate in both wildtype and mutant embryos.

### RNA-sequencing

Ten to twenty manually dechorionated 24 hpf embryos of a germline mutant incross and a germline wildtype incross were homogenized in TRIzol (Ambion, 15596018). Subsequently, the Quick RNA microprep kit (Zymo Research, R1051) was used to isolate RNA and treat the samples with DNAsel. Most samples were depleted from rRNA using the Ribo-Zero rRNA Removal Kit (Illumina, MRZH11124), followed by fragmentation, cDNA synthesis, and libraries were generated using the KAPA Hyper Prep Kit (KAPABiosystems, KK8504). Sequencing libraries were paired-end sequenced (43 bp read-length) on an Illumina NextSeq500 platform. However, two samples per genotype were generated with the TruSeq Stranded Total RNA Library Prep Kit with Ribo-Zero (Illumina, RS-122-2201) and single-end sequenced (50 bp read-length) on an Illumina HiSeq 2500. For wildtype and *MZezh2* mutant embryos, 6 and 7 biological replicates were used, respectively.

### Mass spectrometry

At 24 hpf, 50 embryos were collected, dechorionated, and resuspended by gently pipetting in 500 μl deyolking buffer (1/2 Ginzburg Fish Ringer without Calcium: 55 mM NaCl, 1.8 mM KCl, 1.25 mM NaHCO3, 1x cOmplete EDTA-free protease inhibitor cocktail from Sigma) and incubated for 5 minutes in a Thermomixer at RT at 1,100 rpm to disrupt the yolk. The samples were then centrifuged for 30 seconds at 400 g and the pellet was washed two times in 0.5 ml wash buffer (110 mM NaCl, 3.5 mM KCl, 2.7 mM CaCl2, 10mM Tris/Cl pH8.5, 1x cOmplete EDTA-free protease inhibitor cocktail from Sigma) for 2 minutes in a Thermomixer at RT and 1,100 rpm, followed by 30 seconds centrifugation at 400 g. Washed pellets were lysed in 100 μl RIPA buffer (50 mM Tris pH8.0, 150 mM NaCl, 0.1% SDS, 1% NP-40, 0.5% DOC, 20% glycerol, 1 mM Sodium Orthovanadate, 1x cOmplete EDTA-free protease inhibitor cocktails from Sigma) and sonicated for 2 cycles of 15s ON and 15s OFF on full power at 4°C on a Bioruptor (Diagenode). Samples were incubated for 1 hour on a rotating wheel at 4°C and centrifuged 10 minutes at 12,000 g and 4°C. Supernatant was flash frozen and stored at −80°C. After Bradford analysis, 100 μg protein lysate was used for FASP-SAX as previously described (Wisniewski et al., 2011). The peptide fractions were separated on an Easy nLC 1000 (Thermo Scientific) connected to a Thermo scientific Orbitrap Fusion Tribrid mass spectrometer. MS and MS/MS spectra were recorded in a top speed modus with a run cycle of 3s using Higher-energy Collision Dissociation (HCD) fragmentation. The raw mass spectrometry data were analyzed using the MAXQuant software version 1.6.0.1 (http://www.ncbi.nlm.nih.gov/pubmed/19029910) with default settings. Data was searched against the *Danio rerio* data base (UniProt June 2017). The experiment was performed with biological triplicates for each condition.

### Bioinformatics analyses

For ChIP-seq analysis, fastq files were aligned to GRCz10 zebrafish genome version using BWA-MEM (version 0.7.10-r789) for paired-end reads (Li and Durbin, 2009). Duplicated and multimapping reads were removed using samtools (Li et al., 2009) version 1.2 and Picard tools (http://broadinstitute.github.io/picard) version 2.14.1. When spike-in normalization was used, *Drosophila* reads were aligned to dm6 *Drosophila* genome version and a normalization factor was then applied to zebrafish reads according to manufacturer’s protocol (Active Motif, 53093 and 61686). MACS2 (Zhang et al., 2008) version 2.1.1 was used to call peaks from each aligned bam files using an Input track from 24 hpf wild-type embryos as control sequence. Peaks separated by less than 1kb distance were merged, peaks that were called using Input alone were removed from all data sets using bedtools suit version 2.20.1, and the intersection between the replicates for each antibody in each condition was used to define the definitive peak sets. For visualization in heatmaps and genome browser snapshots, fastq files from duplicate ChIP-sequencing were merged, aligned as described above, subsampled to equalized read numbers between wildtype and *MZezh2* mutant conditions for each ChIP, and transformed into bigwig alignment files using bam2bw version 1.25. Peak lists were analyzed using bedtools and heatmaps were produced using deepTools plotHeatmap (Ramirez et al., 2016) version 2.5.3. Comparison between H3K4me3 peaks in *MZezh2* mutant and wildtype conditions was performed using DiffBind version 2.10.0 on the union between H3K4me3 peaks detected in both conditions.

For RNA-sequencing analysis, read counts per gene were retrieve using GeneCounts quantification method from STAR (Dobin et al., 2013) version 2.4.0 and the GRCz10 zebrafish genome version with Ensembl annotation version 87 as reference. Differential expression analysis was calculated with DESeq2 (Love et al., 2014) version 1.14.1.

For proteomics analysis, differential expression of protein between conditions was assessed with DEP (Zhang et al., 2018) version 1.2.0.

Gene Ontology analyses on selected genes were performed using DAVID bioinformatics resources (Huang da et al., 2009) version 6.8 and anatomical term enrichment was done using ZEOGS (Prykhozhij et al., 2013).

### Whole mount *in situ* hybridization

Embryos at 24 hpf were dechorionated and fixed overnight at 4°C in 4% PFA in PBST (0.1% Tween), after which they were gradually transferred to 100% methanol. Prior to ISH, embryos were gradually transferred back to PBST and, subsequently, ISH was performed as described previously (Houwing et al., 2007). The embryos were imaged by light microscopy on a Leica MZFLIII, equipped with a DFC450 camera.

### RT-qPCR analyses

Total RNA was isolated using Trizol from 20 flash-frozen dechorionated 24 hpf wildtype and *MZezh2* mutant embryos cut in two with tweezers. Reverse transcription was achieved using Superscript III (Invitrogen, 18080093) and poly-dT primers. Standard qPCR using SYBR Green (iQ SYBR Green Supermix, BioRad, 1708880) was performed using the primers shown in Table S2. Relative expression was calculated based on expression of housekeeping genes *β-actin.* Calculations were based on at least 3 independent replicates for both conditions.

## Acknowledgements

We thank J. Bakkers, from the Hubrecht Institute, for providing the *tbx2a, tbx2b, tbx3a, tbx5a,* and *isl1* plasmids and J. den Hertog from the Hubrecht Institute for providing the *gsc* plasmid for ISH probe generation. We thank T. Spanings and A. van der Horst from the Radboud University for excellent zebrafish husbandry and E. Janssen-Megens from the Radboud University for excellent technical support. We thank R. Lindeboom, from the Radboud University, for computational advice. We thank Dr. G.J.C. Veenstra, from the Radboud University, and his team for fruitful discussions. We thank Dr. R. Knight, from the King’s College London, for his help with ISH analysis.

## Competing interests

The authors declare no competing interests.

## Funding

The work was funded by the Innovative Research scheme of the Netherlands Organisation for Scientific research (www.nwo.nl, NWO-Vidi 864.12.009, NWO-Meervoud 836.13.003 L.M.K.), the Radboud University Nijmegen Medical Centre tenure track fellowship (www.radboudumc.nl, L.M.K.), the European Union’s Horizon 2020 research and innovation programme under the Marie Sklodowska-Curie Grant (Agreement No. 705939, K.A.), the Howard Hughes Medical Institute and the Huntsman Cancer Institute core facilities (CA24014, B.R.C.), and the Eunice Kennedy Shriver National Institute of Child Health and Human Development of the NIH (T32HD007491, P.J.M.).

## Data availability

The sequencing data have been submitted to the NCBI Gene Expression Omnibus (GEO; http://www.ncbi.nlm.nih.gov/geo/) under accession number GSE119070. The mass spectrometry proteomics data have been deposited to the ProteomeXchange Consortium via the PRIDE (Vizcaino et al., 2016) partner repository with the dataset identifier PXD010922. Reviewers can obtain access to the datasets via login information provided to the editor.

