## Supplemental Files for "Maintenance of spatial gene expression by Polycomb-mediated repression after formation of a vertebrate body plan"

**Running title: PcG repression in zebrafish**

**Authors:** Julien Rougeot[1,2*], Naomi D. Chrispijn[1], Marco Aben[1,2], Dei M. Elurbe[1,2], Karolina M. Andralojc[1], Patrick J. Murphy[3,4], Pascal W.T.C. Jansen[5], Michiel Vermeulen[5], Bradley R. Cairns[3], Leonie M. Kamminga[1,2,*]

**Affiliations:** [1]Radboud University, Faculty of Science, Department of Molecular Biology, Radboud Institute for Molecular Life Sciences, Nijmegen, the Netherlands [2]Radboud University Medical Center, Department of Molecular Biology, Nijmegen, the Netherlands [3]Howard Hughes Medical Institute, Department of Oncological Sciences and Huntsman Cancer Institute, University of Utah School of Medicine, Salt Lake City, UT 84112, USA [4]Wilmot Cancer Institute, Rochester Center for Biomedical Informatics, University of Rochester Medical Center, Rochester, NY [5]Radboud University, Department of Molecular Biology, Faculty of Science, Radboud Institute for Molecular Life Sciences, Oncode Institute, Nijmegen, The Netherlands. *Corresponding authors.

Key words: Polycomb, Ezh2, zebrafish, ChIP-seq, RNA-seq, proteomics

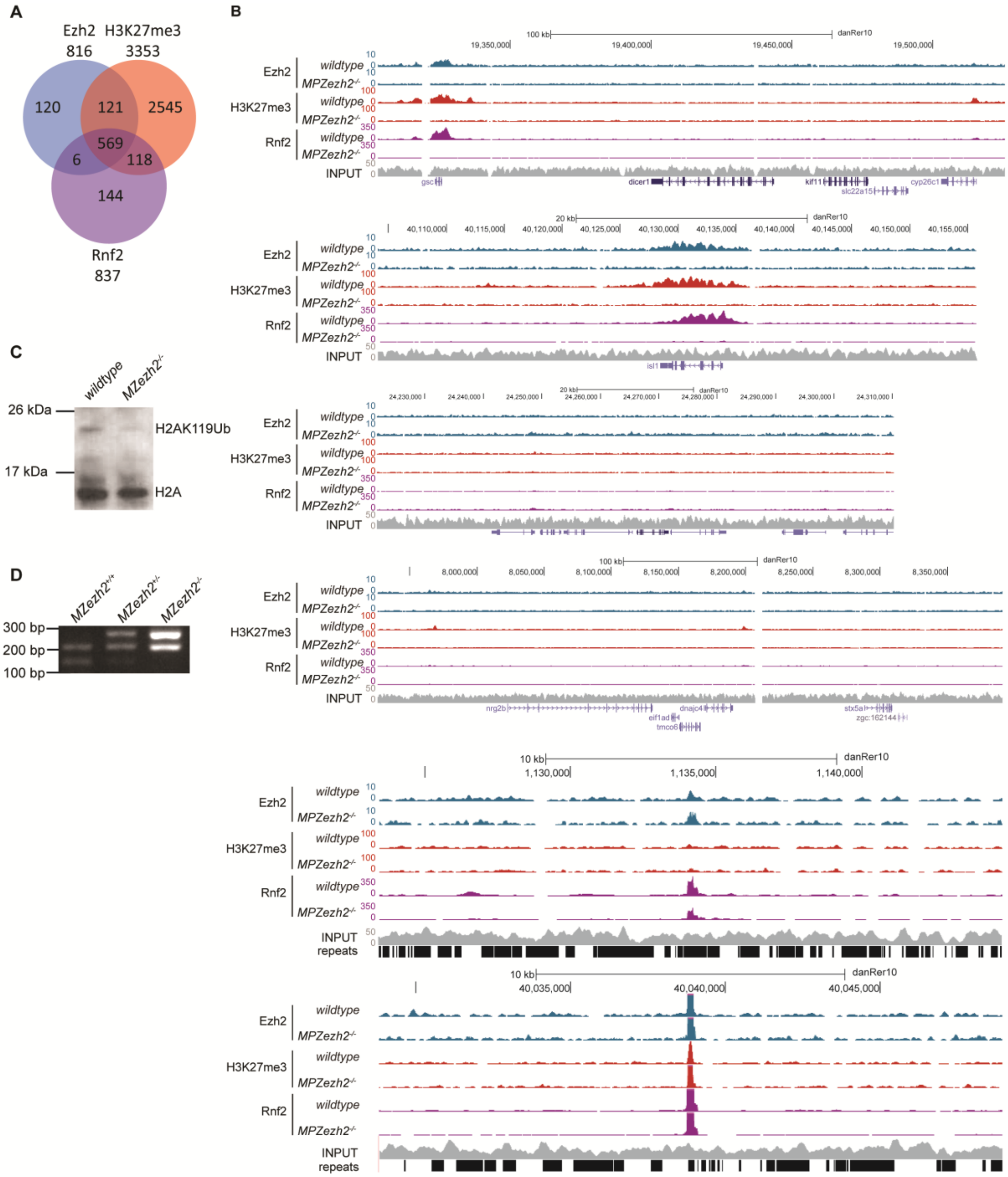


Rougeot_Supplemental_Fig.1

**Supplementary Fig. 1. Analysis of Ezh2, H3K27me3, and Rnf2 binding in wildtype and *MZezh2* mutant (*MZezh2$^{-/-}$*) embryos at 24 hpf.** (**A**) Venn diagrams presenting the overlap between Ezh2 (blue), H3K27me3 (red), and Rnf2 (purple) peaks detected in 24 hpf wildtype embryos. (**B**) UCSC browser snapshots of six genomic loci depicting Ezh2, H3K27me3, and Rnf2 binding after ChIP-seq in *MZezh2$^{-/-}$* embryos compared to wildtype embryos at 24 hpf. Colors represent ChIP-seq for different proteins with blue: Ezh2, red: H3K27me3, purple: Rnf2, and grey: Input control. (**C**) Western blot analysis of H2A on histone extracts at 24 hpf in wildtype and *MZezh2$^{-/-}$* embryos. The presence of H2AK119 monoubiquitylation was visualized as a shift in H2A band from 13 kDa to ≥20 kDa as showed by van der Velden et al. (2012). (**D**) Example of *ezh2$^{(hu5670)}$* genotyping results after nested PCR, RsaI restriction, and gel electrophoresis in *MZezh2* wildtype (*MZezh2$^{+/+}$*), *MZezh2* heterozygous (*MZezh2$^{+/-}$*), and *MZezh2* mutant (*MZezh2$^{-/-}$*) embryos.

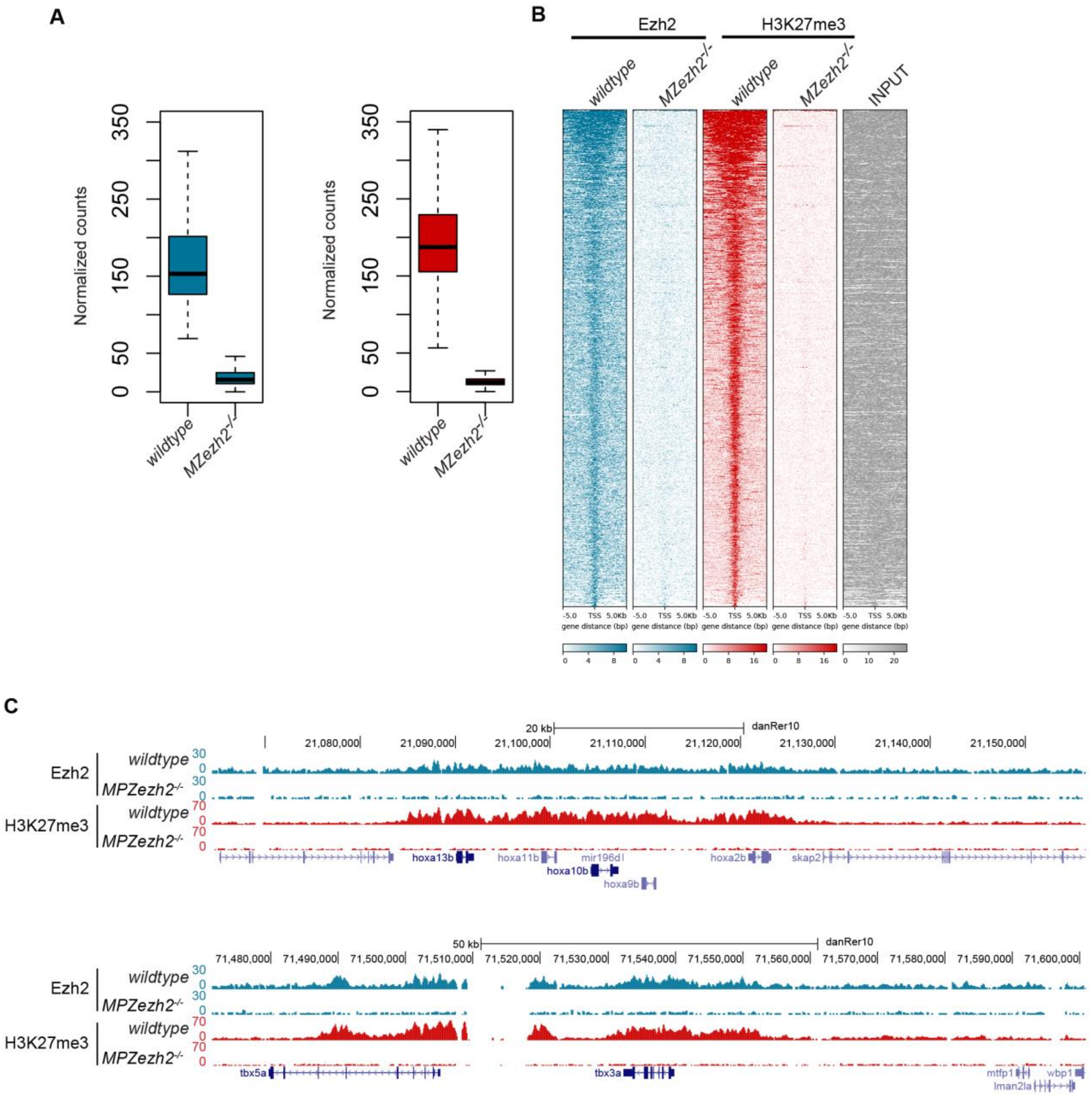


Rougeot_Supplemental_Fig.2

**Supplementary Fig. 2. ChIP-seq of Ezh2 and H3K27me3 using spike-in chromatin for normalization.** (**A**) Box plots of Ezh2, H3K27me3, and Rnf2 coverage based on spike-in normalization after ChIP-seq in wildtype and in *MZezh2*$^{-/-}$ embryos at 24 hpf. Coverages were calculated based on positions of peaks detected in wildtype embryos. (**B**) Heatmaps for Ezh2, H3K27me3, and Rnf2 counts normalized with spike-in chromatin after ChIP-seq in 24 hpf wildtype and *MZezh2*$^{-/-}$ embryos. Windows of 10 kb regions for all H3K27me3 o Ezh2 peaks in 24 hpf wildtype embryos are shown. The input track obtained from 24 hpf wildtype embryos was used as control and was not normalized. (**C**) UCSC genome browser snapshot depicting the loss of Ezh2 and H3K27me3 after ChIP-seq in 24 hpf *MZezh2*$^{-/-}$ embryos compared to wildtype embryos. Coverage were normalized with spike-in chromatin.

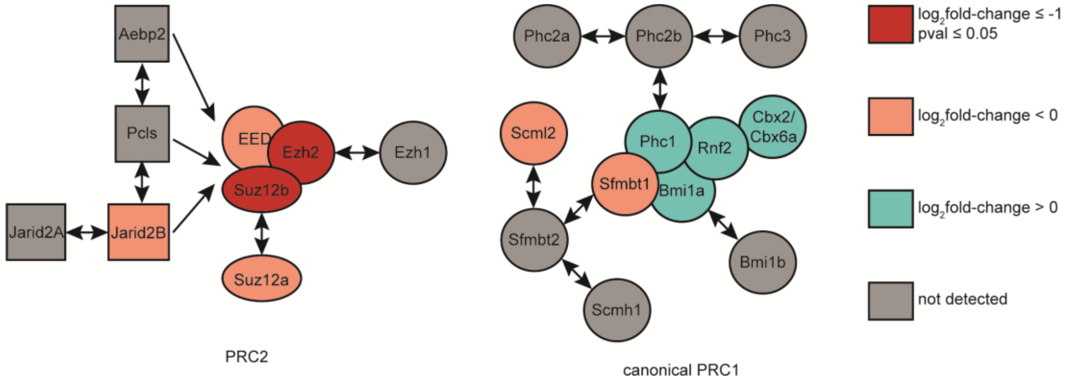


Rougeot_Supplemental_Fig.3

**Supplementary Fig. 3. RNA-seq and proteomics analysis in *MZezh2* mutant (*MZezh2*$^{-/-}$) embryos at 24 hpf.** Schematic representation of changes in protein expression level of PRC2 (left) and canonical PRC1 (right) subunits in *MZezh2*$^{-/-}$ compared to wildtype embryos at 24 hpf. Dark red: $\log_2$fold-change ≥ 1 and *P*-value ≤ 0.05, light red: $\log_2$fold-change ≥ 0, turquoise: $\log_2$fold-change ≤ 0, grey: protein not detected.

**Supplementary Table 2. List of primers used in this study**

| name | sequence | experiment | target |
|---|---|---|---|
| p3_hu5670_ComFw | CAGAATCGGTTTCCAGGTTGCCG | genotyping | *ezh2* genomic PCR |
| p4_hu5670_ComRv | CAGTACTCTGAGATGAACTCATTC | genotyping | *ezh2* genomic PCR |
| LK_ezh2_exon_Fw | TGTAAAACGACGGCCAGTCAGAATCGGTTTCCAGGTTGCCG | genotyping | *ezh2* genomic nested PCR |
| LK_ezh2_exon_Rv | AGGAAACAGCTATGACCATTGCAGGAGACGTTTTTACTGTCCC | genotyping | *ezh2* genomic nested PCR |
| Hoxa9a_RTqPCR_Fw | AAGCAGAATCTAGCCGAACTG | RT-qPCR | *hoxa9a* |
| Hoxa9a_RTqPCR_Rv | CACAGGGTTTTCTGGATCAGC | RT-qPCR | *hoxa9a* |
| Hoxa9b_RTqPCR_Fw | CAACGGATCACATGATGAGAAAAT | RT-qPCR | *hoxa9b* |
| Hoxa9b_RTqPCR_Rv | CCAGTTGGACGAAGGGTTA | RT-qPCR | *hoxa9b* |
| Hoxa11b_RTqPCR_Fw | AGCAGCAATGGACAAAAGACAC | RT-qPCR | *hoxa11b* |
| Hoxa11b_RTqPCR_Rv | AAGAAAAATTCTCTCTCCAGCTCT | RT-qPCR | *hoxa11b* |
| Hoxa13b_RTqPCR_Fw | GTGTACTGCCCGAAAGATCA | RT-qPCR | *hoxa13b* |
| Hoxa13b_RTqPCR_Rv | ACCTGACACGGTATCTTGGA | RT-qPCR | *hoxa13b* |
| tbx2a_RTqPCR_Fw | GCTAAGGAGCTTTGGGATCA | RT-qPCR | *tbx2a* |
| tbx2a_RTqPCR_Rv | CACCTTGAACGGAGGAAACA | RT-qPCR | *tbx2a* |
| tbx2b_RTqPCR_Fw | TCTCAACACATGCTTGCCTC | RT-qPCR | *tbx2b* |
| tbx2b_RTqPCR_Rv | AAAAGTCCACCGAAGGTTGG | RT-qPCR | *tbx2b* |
| tbx3a_RTqPCR_Fw | CCCGATGCCGTTTCATCTG | RT-qPCR | *tbx3a* |
| tbx3a_RTqPCR_Rv | CCGAAAGGAGACATAGCCAG | RT-qPCR | *tbx3a* |
| tbx5a_RTqPCR_Fw | GGGAGCTGATACGAGCTTTT | RT-qPCR | *tbx5a* |
| tbx5a_RTqPCR_Rv | CGTGAGGCCTTAAATTCCGA | RT-qPCR | *tbx5a* |
| isl1_RTqPCR_Fw | TTTACAAATGGCAGCAGAGC | RT-qPCR | *isl1* |
| isl1_RTqPCR_Rv | CGGGTTGTTTTCTCAGGTTG | RT-qPCR | *isl1* |
| gsc_RTqPCR_Fw | CAACAGTGTCCGTGTATTCCT | RT-qPCR | *gsc* |
| gsc_RTqPCR_Rv | TCATTTGATGTGGGACTGGAG | RT-qPCR | *gsc* |

**Supplemental Table S3. List of antibodies used in this study**

| antibody | brand | ref | Concentration µg/µl | ChIP (µl/IP) | WB (dilution) |
|---|---|---|---|---|---|
| anti-Ezh2 | Cell Signaling | 5246S | N/A | 2 | 1:1,000 |
| anti-Rnf2 | Cell Signaling | 5694S | N/A | 4 | N/A |
| anti-H3K27me3 | Millipore | 07-449 | N/A | 2 | N/A |
| anti-H3K4me3 | Millipore | 04-745 | N/A | 2 | N/A |
| anti-H2A | Millipore | 07-146 | N/A | N/A | 1:1,000 |
| anti-Histone H3 | Sigma-Aldrich | H0164 | N/A | N/A | 1:2,000 |
| HRP-conjugated anti-Rabit | Dako | P0217 | N/A | N/A | 1:3,000 |
